# Nucleic acid 3D structure search and alignment with GTalign

**DOI:** 10.64898/2026.06.01.729228

**Authors:** Mindaugas Margelevičius, Shikhar Rana

## Abstract

**Summary:** Structural comparison of nucleic acids, particularly RNA, is critical for understanding evolutionary and functional relationships beyond sequence similarity, yet efficient tools for large-scale 3D structure search and alignment remain scarce. We extend GTalign to support nucleic acid structures, enabling unified, high-performance alignment across macromolecules. Benchmarking on diverse nucleic acid datasets demonstrates improved alignment accuracy and substantially lower runtimes compared to existing methods. GTalign thus provides a scalable solution for nucleic acid structure comparison and database search. Availability and Implementation: https://github.com/minmarg/gtalign_alpha

## Introduction

Structural comparison of macromolecules is a fundamental task in computational structural biology, which enables the detection of evolutionary relationships, functional similarities, and structural motifs that are often not evident from sequence alone. While numerous algorithms have been developed for comparison of protein three-dimensional (3D) structures, efficient tools for large-scale search and alignment of nucleic acid structures remain scarce.

Nucleic acids, in particular RNAs, often exhibit complex tertiary folds and structural motifs that are conserved despite substantial sequence divergence (Boerneke et al., 2014), making structural comparison a natural approach for their analysis.

The problem of RNA 3D structure alignment has been addressed using a variety of algorithmic strategies. Early approaches focused on identifying conserved structural regions, such as secondary structure units or base-pairing properties, to guide alignment and improve both accuracy and efficiency (Ferrè et al., 2007; Hoksza and Svozil, 2012; Ge and Zhang, 2015). To reduce the complexity of direct 3D comparison, some methods transform structures into one-dimensional representations (Yang et al., 2016) or apply sequence alignment techniques based on structural similarity (Capriotti and Marti-Renom, 2008). Other approaches employ seed-and-extend strategies, identifying local structural anchors using hashing schemes and expanding them into full alignments (Dror et al., 2005; Bauer et al., 2009). Related methods detect small sets of spatially proximal atoms (e.g., cliques or triplet seeds) and extend these using either greedy (Rahrig et al., 2010; Nguyen et al., 2017; Piątkowski et al., 2017) or combinatorial search procedures (Zurkowski et al., 2023, 2024; Bohdan et al., 2024).

Among widely used tools, RNA-align (Gong et al., 2019), now incorporated into US-align (Zhang et al., 2022), applies iterative superposition of structural fragments to construct alignments and achieves improved runtime compared to earlier methods. Nevertheless, scalable approaches for efficient search and alignment of nucleic acid 3D structures across large databases remain needed.

Here we present an extended version of GTalign (Margelevičius, 2024), a high-performance structure alignment method originally developed for fast and accurate alignment of protein 3D structures. The new implementation expands its capabilities to support nucleic acid structures (Supplementary Section S1.1), enabling unified, high-performance search and alignment across both protein and nucleic acid structure datasets while maintaining the speed and accuracy of the original framework. We demonstrate that GTalign achieves higher alignment accuracy and substantially faster runtimes compared to existing methods on large-scale nucleic acid structure benchmarks.

## Materials and methods

We benchmarked GTalign against US-align, ARTEMIS (Bohdan et al., 2024), RMalign (Zheng et al., 2019), and RTM-align (Qiu et al., 2024) (Supplementary Section S1.2). These methods use different strategies for deriving structural alignments. RMalign and RTM-align follow an optimization strategy similar to RNA-align but modify the TM-score (Gong et al., 2019). In particular, they modify the scaling factor (*d*_0_) to improve sensitivity for short RNA structures. RTM-align additionally applies score standardization and a sigmoid transformation. ARTEMIS uses its own alignment method and evaluates the resulting alignments using the original TM-score formulation.

Given the widespread use of US-align and TM-score in nucleic acid structure comparison, this group of TM-score-based and recent methods provides a representative benchmark for comparison. Other methods with very long runtimes (days to weeks) on the benchmark dataset were excluded, as were tools that were unavailable or could not be compiled.

## Datasets

Two datasets were used for benchmarking. The first was the non-redundant set of 637 RNA structures obtained from the US-align RNA dataset (Zhang et al., 2022). An all-against-all comparison was performed, excluding self-matches. The structures range in length from 30 to 4449 nucleotides and share less than 80% sequence identity. The second dataset comprised the first nucleic acid chain of each such PDB entry with a length of at least 20 nucleotides, resulting in 5025 chains in total. Structurally diverse query structures were obtained by clustering this dataset at a TM-score threshold of 0.4 and a minimum coverage of 60%, followed by selecting the 40 longest members shorter than 4000 nucleotides from singleton clusters (Supplementary Section S1.3). Of these queries, 10 represented DNA chains.

### Evaluation metrics

Alignment accuracy was evaluated using root-mean-square deviation (RMSD) of aligned nucleotides and TM-score (Gong et al., 2019), which capture local and global structural similarity, respectively. Higher TM-scores for a given pair of structures generally indicate more accurate alignments, with a value of 1 corresponding to a perfect match.

TM-score can be normalized by the length of either structure. Here, we report the higher value obtained for each pair (i.e., normalized by the shorter structure), although very similar results were observed when normalizing by query length.

TM-score and RMSD for a given alignment were computed using the TM-align program (Zhang and Skolnick, 2005), with minor modifications to support nucleic acid structures. Values reported by US-align were used as provided. (Independent verification confirmed that they agree with values obtained using the modified TM-align.) The evaluation procedure followed protocols previously applied to protein structure alignment (Margelevičius, 2024) and to the alignment of macromolecular complexes, including RNA-containing complexes (Margelevičius, 2025).

Runtimes were measured using the Linux `time` command.

### System configuration

Benchmarking of GTalign and US-align was performed on a server with two Intel Xeon Gold 5115 CPUs (2.4 GHz; 20 hardware threads per CPU), 128 GB DDR4 RAM, and three NVIDIA V100-PCIE-16GB GPUs. The GPU implementation of GTalign used a single GPU, while US-align was executed using all 40 CPU cores. The remaining methods were run on a cluster node with 80 CPU cores (Intel Xeon Gold 6148, 2.4 GHz) to mitigate their substantially longer runtimes. Runtime comparisons with these methods were not intended to directly compare their performance with GTalign. Rather, the primary runtime comparison was between GTalign and US-align, the two fastest methods in our benchmark.

## Results and discussion

Benchmark results are shown in Figure 1. For the second dataset, only US-align, its fast variant, and GTalign were evaluated, as the other tools either do not support the mmCIF format (RMalign) or required prohibitively long runtimes.

**Figure 1.**
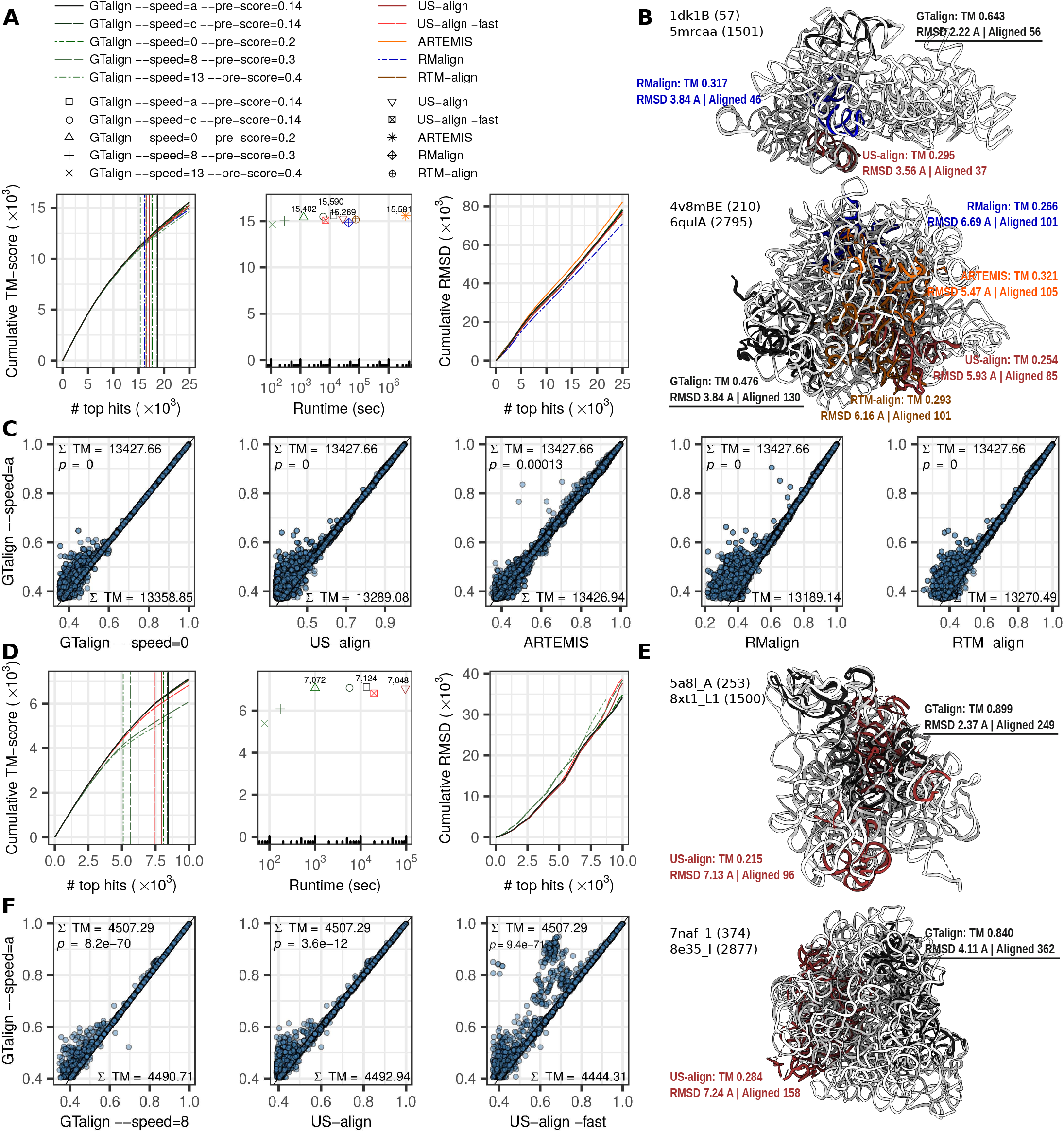
Benchmark results on the RNA (Zhang et al., 2022) (A–C) and PDB-derived (D–F) datasets. (A,D) Left: cumulative TM-score as a function of the number of top-ranked alignments. Vertical lines indicate the number of alignments with TM-score*≥*0.45. Middle: cumulative TM-score as a function of runtime (seconds). Right: cumulative RMSD as a function of the number of top-ranked alignments, ranked by each tool’s reported similarity score. (B,E) Representative rRNA alignment examples. Structure lengths are given in parentheses. The larger structure is shown in gray, while the smaller structures are colored and superimposed according to each tool. ARTEMIS and RTM-align alignments in (B) (upper panel) are nearly identical to those of GTalign and US-align, respectively, and are not shown. (C,F) Scatter plots of TM-scores for the top 20,516 (C) and 6,004 (F) alignments identified in common by all tools. Σ TM denotes the cumulative TM-score over these alignments for each tool. TM, TM-score; *p, p*-values obtained using the *paired* two-sided Wilcoxon signed-rank test; *p* = 0, values below 2.2 *×* 10^−16^.

We evaluated several GTalign parameterizations to characterize its performance. The `--speed` option controls the exploration depth of the superposition space (range a–c and 0–13), with letters and lower values corresponding to more exhaustive search. The `--pre-score` option enables similarity prescreening based on provisional TM-scores. The default `--pre-score` value is 0.4 for proteins, whereas lower values (0.2–0.3) are recommended for RNAs due to its greater structural flexibility. Consistently, while a TM-score of at least 0.5 typically indicates similar topology for proteins (Xu and Zhang, 2010), the corresponding threshold for RNAs is 0.45 (Gong et al., 2019). Results are reported for `--speed={a,c,0,8,13}` and `--pre-score={0.14,0.2,0.3,0.4}`.

The left panels of Figure 1A,D show cumulative TM-score as a function of the number of top-ranked alignments. For the first dataset (Fig. 1A), the most exhaustive GTalign configuration (`--speed=a`) achieved the highest cumulative TM-score (15,590), followed by ARTEMIS (15,581), and identified more significant matches (TM-score ≥ 0.45; 18,779 vs. 18,696). For the second dataset (Fig. 1D), GTalign (`--speed=a`) similarly outperformed US-align in cumulative TM-score (7,124 vs. 7,048), while identifying approximately 6% more significant matches (8,449 vs. 7,970).

The advantage of GTalign is also evident among the 20,516 (Fig. 1C) and 6,004 (Fig. 1F) top hits identified in common. The corresponding TM-score scatter plots show that GTalign yields, on average, higher scores for the same alignments.

The cumulative RMSD distributions are broadly similar across methods, except for ARTEMIS and RMalign in the first dataset (Fig. 1A, right) and the fastest GTalign configurations in the second dataset (Fig. 1D, right). ARTEMIS consistently produces higher RMSD values despite generating alignments of similar average length (502 nucleotides) to GTalign (499) and US-align (500) (Supplementary Table S1). In contrast, RMalign produces lower RMSDs, which can be attributed to its shorter alignments (488 nucleotides on average). Shorter alignments generally reduce RMSD but may also reduce TM-score. Thus, RMSD alone does not necessarily indicate better structural agreement.

Figure 1B,E shows two representative ribosomal RNA alignments for the respective datasets. In these examples, GTalign identifies more accurate alignments at distinct structural locations compared to other methods, resulting in lower RMSDs and higher TM-scores. While such cases are not universal, the overall tendency of GTalign to produce higher TM-scores reflects its ability to identify more accurate structural matches (Fig. 1A,D).

GTalign achieves this accuracy at substantially lower computational cost (Fig. 1A,D, middle). At an exhaustive setting (`--speed=0`), GTalign was 22x and 95x faster than US-align (1,287 s vs. 27,736 s and 1,004 s vs. 95,646 s, respectively), while achieving higher cumulative TM-scores (15,402 vs. 15,269 and 7,072 vs. 7,048, respectively). The fastest configuration (`--speed=13`, `--pre-score=0.4`) was 66x and 251x faster than US-align -fast (111 s vs. 7,336 s and 78 s vs. 19,612 s, respectively), while retaining most of the accuracy of `--speed=0` (Fig. 1A).

This efficiency is achieved on an older-generation GPU. GTalign also provides a CPU implementation for broader applicability, with corresponding runtimes reported in Supplementary Table S2. Although the CPU implementation is >10x slower than the GPU implementation, the GTalign algorithm was specifically designed and optimized for GPU execution, providing substantial advantages in computational speed and energy efficiency, with CPU execution requiring approximately an order of magnitude more power.

## Conclusion

We presented an extension of GTalign for efficient search and alignment of nucleic acid 3D structures. Benchmarking on diverse datasets showed that GTalign achieves higher alignment accuracy than existing methods, with higher TM-scores and more significant matches, while substantially reducing computational cost. The resulting method provides a practical and scalable approach for nucleic acid structure comparison and database search.

The updated functionality is also available through the GTalign webserver (Dapkūnas and Margelevičius, 2025) (https://bioinformatics.lt/comer/gtalign), enabling unified, high-throughput searches across protein and nucleic acid structure databases. This capability facilitates large-scale structural analyses and the identification of conserved structural motifs and functional relationships across diverse macromolecules.

## Data availability

The GTalign source code is available at https://github.com/minmarg/gtalign_alpha and archived on Zenodo (version 1.1.0; https://doi.org/10.5281/zenodo.22871255). Dataset information, benchmark data, and scripts for benchmarking and figure generation are available at https://github.com/minmarg/gtalign-rna-evaluation. The 637-structure US-align RNA dataset (Zhang et al., 2022) is available at https://doi.org/10.6084/m9.figshare.16725745.

## Supplementary Information

### S1 Supplementary methods

#### S1.1 Algorithm extension to nucleic acids

GTalign was originally developed for searching and aligning protein structures [1]. Its core algorithm is based on spatial indexing of structures and the use of these indices to rapidly identify favorable structural superpositions. In principle, this framework is applicable to macromolecular structures of different types. However, several components of the original implementation were specific to proteins. For example, spatial indices are applied twice for each structure pair, with one of the applications incorporating secondary structure information. In addition, protein-specific TM-scores [2] were used throughout the alignment process.

To support nucleic acids and provide a unified framework for heterogeneous monomer analysis, GTalign was extended in several respects: (i) support for multiple atom types and molecular categorization; (ii) secondary structure assignment for nucleic acids; (iii) nucleic acid-specific structural similarity metrics; (iv) a revised strategy for generating initial superpositions; (v) modified similarity pre-screening; (vi) deeper exploration of the superposition space; and (vii) high-performance GPU (CUDA) and CPU (OpenMP) parallel implementations.

The nucleic acid-specific similarity metrics are based on the RNA-specific TM-score formulation. GTalign calculates these scores consistently with the established formulation [3]. The TM-score formula [2] is identical for proteins and RNAs, 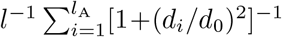, where *l* is the length of the structure used for normalization (the shorter of the two structures in our benchmarks), *l*_A_ is the alignment length, *d*_*i*_ is the distance between corresponding spatially aligned residues, and *d*_0_ is a length-dependent scaling factor. The difference between the protein and RNA formulations lies in *d*_0_. For RNA structures, *d*_0_ is calculated as 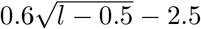 for *l* ≥ 30 and is assigned a constant value smaller than 1 for shorter structures [3, 4]. GTalign allows the representative atoms used for TM-score calculation to be specified flexibly. By default, nucleotides are represented by their C3’ atoms.

The revised seeding strategy for initial superpositions was recently introduced in GTcomplex [5] for macromolecular complex alignment and has now been adapted for GTalign. The effectiveness of identifying an optimal superposition depends strongly on how initial seeds representing candidate superpositions are generated [1]. In the new strategy, GTalign generates local alignment seeds based on local structural similarity. This approach improves the efficiency of initial superposition estimation and generally leads to more accurate final alignments. The local alignment-based seeding strategy is now used by default for both nucleic acid and protein structures.

Similarity pre-screening based on provisional TM-scores works effectively for proteins. For nucleic acids, however, provisional scores correlate less strongly with the final alignment scores. Consequently, achieving high sensitivity requires a lower similarity pre-screening threshold (`--pre-score`), as demonstrated by the benchmark results, at the cost of reduced computational efficiency.

Although provisional scores are calculated and applied separately for proteins and nucleic acids, the pre-screening strategy is not yet fully optimized for their distinct structural characteristics. For example, the strict incorporation of secondary structure information into spatial indexing may be less beneficial for nucleic acids because of their greater structural flexibility. This and other aspects will be investigated in future work to improve similarity pre-screening for nucleic acid structures.

#### S1.2 Command-line configurations of benchmarked tools

The command-line configurations used for benchmarking are described below. US-align, RMalign, and RTM-align searches were parallelized across CPU cores by distributing queries among available workers.

GTalign version 1.1.0 was used with the following command:

~~~
gtalign --qrs=<query_dir> --rfs=<db_dir> -o <output_dir> -c <cache_dir> --add-search-by-
   ss --speed=<speed> --pre-score=<prescore_threshold> -s 0.33 --dev-queries-total-
   length-per-chunk=<qlength> --dev-max-length=<slength> --cpu-threads-reading=2
~~~

where `<query_dir>` and `<db_dir>` specify the directories containing query and subject structures, respectively; `<output_dir>` specifies the output directory; and `<cache_dir>` specifies a temporary directory for cached data. `<speed>` and `<prescore_threshold>` denote the evaluated `--speed` configurations and similarity-prescreening cutoff, respectively. The total length of queries processed as a single chunk, `<qlength>`, was set to 4500 for the RNA dataset [4] and left at its default value for the PDB-derived dataset. The maximum subject length, `<slength>`, was set to 4500 and 8100 for the RNA and PDB-derived datasets, respectively.

US-align [4] version 20241108 was executed using:

USalign `<query_file>` -dir2 `<db_dir>` `<lst_file>` [-fast]

where `<query_file>` specifies the query structure, `<db_dir>` the subject directory, and `<lst_file>` a list of subject structures. The -fast option was used for the fast configuration.

ARTEMIS [6] version 1.51 was executed using:

~~~
artemis.py -q <query_dir> -r <db_dir> -s -threads 80
~~~

RMalign [7] version 1.1 was executed using:

~~~
RNAalign -A <query_file> -B <subject_file>
~~~

where `<query_file>` and `<subject_file>` specify query and subject structures, respectively.

RTM-align [8] was executed using:

~~~
RTMalign <query_file> <subject_file>
~~~

#### S1.3 Selection of structurally diverse nucleic acid queries

One of the two benchmark datasets comprised all 5025 first nucleic acid chains from the PDB [9] (dated 4/5/2026) with a length of at least 20 nucleotides. To select structurally diverse query structures, the chains were clustered using GTalign (v1.0.1) with a TM-score threshold of 0.4 and a minimum coverage of 60%, using the command:

~~~
 gtalign -v --cls=mmCIF_na_1st_chains -o <output_dir> --dev-queries-total-length-per-
   chunk=8100 --dev-max-length=8100 --pre-score=0.3 --speed=10 --add-search-by-ss –cls
   -coverage=0.6 --cls-threshold=0.4 -c <cache_dir>
~~~

where `mmCIF_na_1st_chains` specifies the directory containing the corresponding input structures.

The final query set was obtained by selecting the 40 longest structures shorter than 4000 nucleotides from singleton clusters. The query structures range in length from 210 to 2891 nucleotides. Dataset information, including the complete list of queries and the 10 DNA chains among them, is publicly available.

### S2 Supplementary results

#### S2.1 Supplementary tables

**Table S1.** Number of aligned nucleotides and sequence identity obtained by each method.

| | $\bar{n}_{\text{aln}}$ | $\sigma_{n_{\text{aln}}}$ | $\bar{n}_{\text{idn}}$ | $\sigma_{n_{\text{idn}}}$ | $\bar{I}_{\text{aln}}$ | $\bar{I}_{\text{len}}$ |
| --- | --- | --- | --- | --- | --- | --- |
| *RNA dataset [4]* |  |  |  |  |  |  |
| GTalign --speed=a --pre-score=0.14 | 499.415 | 826.353 | 288.949 | 530.584 | 44.80 | 39.88 |
| GTalign --speed=c --pre-score=0.14 | 499.602 | 826.321 | 289.028 | 530.570 | 44.85 | 39.87 |
| GTalign --speed=0 --pre-score=0.2 | 496.566 | 825.856 | 287.596 | 530.473 | 44.84 | 39.67 |
| GTalign --speed=8 --pre-score=0.3 | 485.311 | 826.194 | 282.074 | 531.103 | 44.33 | 38.97 |
| GTalign --speed=13 --pre-score=0.4 | 481.337 | 827.970 | 280.623 | 531.630 | 44.31 | 38.31 |
| US-align | 499.881 | 823.339 | 290.924 | 532.178 | 45.08 | 39.91 |
| US-align -fast | 496.910 | 824.743 | 289.970 | 532.633 | 44.96 | 39.61 |
| ARTEMIS | 502.439 | 827.078 | 289.017 | 528.643 | 44.75 | 40.16 |
| RMalign | 487.648 | 826.901 | 285.600 | 533.378 | 44.42 | 40.29 |
| RTM-align | 498.670 | 821.399 | 289.897 | 530.496 | 45.02 | 39.85 |
| *PDB-derived dataset* |  |  |  |  |  |  |
| GTalign --speed=a --pre-score=0.14 | 466.433 | 430.490 | 274.134 | 302.474 | 51.52 | 44.38 |
| GTalign --speed=c --pre-score=0.14 | 468.397 | 429.371 | 274.838 | 302.005 | 51.60 | 44.35 |
| GTalign --speed=0 --pre-score=0.2 | 465.784 | 428.279 | 271.404 | 301.689 | 51.18 | 44.00 |
| GTalign --speed=8 --pre-score=0.3 | 428.472 | 477.117 | 237.005 | 311.663 | 44.72 | 36.59 |
| GTalign --speed=13 --pre-score=0.4 | 477.155 | 501.328 | 264.745 | 324.135 | 46.22 | 37.75 |
| US-align | 520.436 | 443.163 | 292.097 | 299.272 | 51.67 | 43.71 |
| US-align -fast | 527.225 | 458.788 | 290.443 | 301.823 | 50.18 | 42.15 |

The 25,000 and 10,000 top-ranked alignments were considered for the RNA and PDB-derived datasets, respectively. $\bar{n}_{\text{aln}}$ , mean number of aligned nucleotides; $\sigma_{n_{\text{aln}}}$ , standard deviation of the number of aligned nucleotides; $\bar{n}_{\text{idn}}$ , mean number of aligned identical nucleotides; $\sigma_{n_{\text{idn}}}$ , standard deviation of the number of aligned identical nucleotides; $\bar{I}_{\text{aln}}$ , mean sequence identity percentage calculated over aligned nucleotides; $\bar{I}_{\text{len}}$ , mean sequence identity percentage calculated relative to the length of the shorter structure.

**Table S2.** Cumulative TM-scores (Σ TM-score) and runtimes (seconds) obtained by each method.

| | $\Sigma$ TM-score | $t_{\text{GPU}}$ (s) | $t_{\text{CPU}}$ (s) |
| --- | --- | --- | --- |
| *RNA dataset* |  |  |  |
| GTalign --speed=a --pre-score=0.14 | 15,590 | 13,494 | 205,568 |
| GTalign --speed=c --pre-score=0.14 | 15,563 | 5,789 | 83,344 |
| GTalign --speed=0 --pre-score=0.2 | 15,402 | 1,287 | 15,867 |
| GTalign --speed=8 --pre-score=0.3 | 15,039 | 289 | 3,182 |
| GTalign --speed=13 --pre-score=0.4 | 14,632 | 111 | 1,184 |
| US-align | 15,269 | – | 27,736 |
| US-align -fast | 15,092 | – | 7,336 |
| ARTEMIS | 15,581 | – | 3,790,159 |
| RMalign | 14,835 | – | 44,020 |
| RTM-align | 15,252 | – | 75,016 |
| *PDB-derived dataset* |  |  |  |
| GTalign --speed=a --pre-score=0.14 | 7,124 | 13,466 | 224,318 |
| GTalign --speed=c --pre-score=0.14 | 7,118 | 5,610 | 88,998 |
| GTalign --speed=0 --pre-score=0.2 | 7,072 | 1,004 | 13,832 |
| GTalign --speed=8 --pre-score=0.3 | 6,082 | 173 | 2,066 |
| GTalign --speed=13 --pre-score=0.4 | 5,394 | 78 | 786 |
| US-align | 7,048 | – | 95,646 |
| US-align -fast | 6,827 | – | 19,612 |

For cumulative TM-scores, the 25,000 and 10,000 top-ranked alignments were considered for the RNA and PDB-derived datasets, respectively. $t_{\text{GPU}}$ , runtime of GTalign on an NVIDIA V100 GPU; $t_{\text{CPU}}$ , runtime on multiple CPU cores. The CPU version of GTalign was run on the server with 40 hardware threads described in the main text (System configuration), using one thread for data reading (`--cpu-threads-reading=1`).

